# Prioritising the eradication of invasive species from island archipelagos with high reinvasion risk

**DOI:** 10.1101/2021.11.04.467372

**Authors:** Viney Kumar, Andre Nunez, Kaitlyn Brown, Kanupriya Agarwal, Samuel Hall, Michael Bode

## Abstract

Eradicating invasive species from islands is a proven method for safeguarding threatened and endangered species from extinction. Island eradications can deliver lasting benefits, but require large up-front expenditure of limited conservation resources. The choice of islands must therefore be prioritised. Numerous tools have been developed to prioritise island eradications, but none fully account for the risk of those eradicated species later returning to the island: reinvasion. In this paper, we develop a prioritisation method for island eradications that accounts for the complexity of the reinvasion process. By merging spatially-explicit metapopulation modelling with stochastic dynamic optimisation techniques, we construct a decision-support tool that optimises conservation outcomes in the presence of reinvasion risk. We applied this tool to two different case studies – rat (*Rattus rattus*) invasions in the Seaforth archipelago in New Zealand, and cane toad (*Rhinella marina*) invasions in the Dampier archipelago in Australia – to illustrate how state-dependent optimal policies can maximise expected conservation gains. In both case studies, incorporating reinvasion risk dramatically altered the optimal order of island eradications, and improved the potential conservation benefits. The increase in benefits was larger in Dampier than Seaforth (42% improvement versus 6%), as a consequence of both the characteristics of the invasive species, and the arrangement of the islands. Our results illustrate the potential consequences of ignoring reinvasion risk, and demonstrate that including reinvasion in eradication prioritisation can dramatically improve conservation outcomes.

## 1 Introduction

One of the most effective and reliable management actions for threatened species conservation is the eradication of invasive species from islands (Jones et al., 2016). Islands contain a disproportionate number of threatened species, which are frequently endangered by the presence of predatory invasive species, such as rodents and cats (Ricketts et al., 2005; Bellard et al., 2016). By removing these threats, island eradications have had global success in saving species from extinction (Holmes et al., 2015; Legge et al., 2018; Brooke et al., 2018; Jones et al., 2016). Island eradications have been repeated and documented so frequently that we can predict their costs, estimate their probability of success, and forecast their likely benefits and side-effects (Baker and Bode, 2021; Holmes et al., 2015).

While eradications deliver conservation benefits, they require large up-front expenditure (Martins et al., 2006; Holmes et al., 2015). They require specialised equipment and personnel, pose considerable logistical obstacles, and face a substantial risk of failure (Russell and Broome, 2016). Given that conservation resources are severely constrained, it is therefore important that island eradication choices be prioritised. Those choices include the islands that are considered (Brooke et al., 2007), the species that are targeted (Helmstedt et al., 2016), and the eradication methods that are employed (Baker et al., 2017). There is now an extensive literature on the prioritisation of island eradication decisions, and a very large number of tools have been created to support those decisions (Baker and Bode, 2021). These tools identify the best islands and invasive species to target for eradication by considering the intrinsic qualities of the islands. The factors considered include the costs of the eradication program, the threat status of the endemic species, the probability of eradication success, and the presence of complicating factors such as human communities.

A critical issue in island eradications is the possibility that the invasive species will reinvade the island after an eradication project is successful (Pichlmueller et al., 2020; Russell et al., 2010). Reinvasion is common on islands near the mainland, and also within archipelagos where nearby islands can act as sources. New Zealand’s Hauraki Gulf, for example, has 16 documented cases of rodent reinvasion (Holmes et al., 2015) from source populations on both the mainland and nearby islands. The Database of Island Invasive Species Eradications (DIISE) (Holmes et al., 2015) has documented 218 successful eradications across 164 different islands that were followed by reinvasion. In other words, 21% of the 1038 eradications in the database were reinvaded. Rodent reinvasions were the most common (186 cases), but reinvasions of birds, cats, ungulates and amphibians have also been observed.

Reinvasion challenges a key advantage of island eradications: the one-off nature of the expenditure. Invasive species management projects on the mainland require the type of consistent funding that conservation organisations find hard to secure (McBRIDE et al., 2007). By contrast, an island eradication is a single, up-front expense. While an island eradication project will cost more than a mainland control project, the additional initial expenditure is justified by the ongoing benefits that it creates in the years that follow. However, if reinvasion becomes a concern, the invasive-free islands must be consistently monitored, and occasionally re-eradicated. From this perspective, the island eradication project then begins to resembles a mainland control project, but in a challenging and expensive location.

While reinvasion risk is acknowledged as an important consideration when choosing island eradication projects, it rarely features in prioritisation schemes. The two exceptions are (Harris et al., 2012; Dawson et al., 2014), who simply recommend that islands with a high reinvasion risk (specifically, those closer to the mainland than the invasive species’ estimated maximum dispersal distance) be avoided, and that groups of islands be targeted simultaneously as single “eradication units”. Neither option treats reinvasion risk with the care that is applied to factors such as the cost of eradication projects, or the threat status of native species. Nor do these approaches explicitly acknowledge that an island’s reinvasion risk can change dynamically, as invasive species are removed from, or invade, nearby islands.

In contrast with island eradication theory, reinvasion (or recolonisation) dynamics are a central element of metapopulation theory and network theory. When applied to conservation problems, these theories treat reinvasion as a state-dependent probability, defined by the invasion state of the surrounding habitat patches (in this case, islands). However, these ideas from metapopulation and network theory have not been incorporated into island eradication prioritisation. At the same time, metapopulation and network management theory is not well-suited to prioritising island eradication projects. These methods tend to downplay the unique characteristics of individual islands including their size, eradication costs, and their threatened species cohorts, instead treating them as identical nodes in a large network (Ross and Pollett, 2010; Sanchirico et al., 2010; Chadès et al., 2011).

In this paper, we formulate a prioritisation method for island eradications that treats reinvasion as a dynamic and stochastic process. The method is a hybrid of island prioritisation and metapopulation theory. As with traditional island eradication prioritisation, our method considers the individual attributes of different islands, such as the conservation benefits they offer and the cost of eradication (Baker and Bode, 2021). However, in accordance with metapopulation theory, our method also focuses on the complex network of connections among islands and mainland, as well as the state-dependent nature of reinvasion risk (Chadès et al., 2011; Southwell et al., 2017). Our approach uses both sets of factors to produce an adaptive, state-dependent island eradication policy, which recommends a target island for prioritisation given the current invasion state of the archipelago.

## 2 Methods

Our methods combine metapopulation modelling and dynamic optimisation into a general framework for prioritising the eradication of invasive species from islands in the presence of substantial reinvasion risk. In particular, we use stochastic patch occupancy metapopulation modelling (SPOM) to describe the reinvasion dynamics, and stochastic dynamic programming (SDP) to identify time- and state-dependent optimal eradication policies. The code needed to apply these methods, as well as their application to two case-studies, are provided in an online repository.

### 2.1 Modelling metapopulation dynamics

The key processes in spatially explicit metapopulation theory (Hanski, 1994) each have analogues in invasive eradication planning for archipelagos. The invasion state of the islands (i.e., which subset of islands is currently inhabited by the invasive species) is captured by the occupancy state of the metapopulation, colonisation dynamics determine the process of reinvasion, and local extinction captures both natural extinctions and eradications.

While there are many types of spatially explicit metapopulation models (Hanski and Gaggiotti, 2004), we choose to model the spatial and temporal dynamics of invaded archipelagos using a stochastic patch occupancy metapopulation model (SPOM), implemented as a discrete-time, finite-space Markov chain. SPOMs focus only on the presence or absence of the invasive species on each island (Possingham et al., 1993), and assume that the three central processes- natural extinction, reinvasion, and eradication - occur sequentially in discrete annual time steps. For each island *a*, there is a probability *e_a_* (albeit a small one) that the population of the invasive species will become naturally extinct in each timestep. If the managers undertake an eradication project on island *a*, then the probability that they will be successful is *d_a_*. Finally, if invasives currently occupy island *a*, then there is a probability *p_ab_* each year that this population will colonise (or recolonise) an unoccupied island *b*. Similarly, for some islands there is also a possibility that a nearby mainland population could invade island *b*, with probability *p_mb_*.

To describe the invasive species metapopulation dynamics as a Markov process, we must define the possible states of the archipelago, and the probability of transitioning between each state. At any given time, an archipelago of *n* islands can only be in one of 2^*n*^ distinct invasion states. We denote the *i^th^* state of the archipelago as *S_i_*, a binary vector, whose *m^th^* component [*S_i_*]_*m*_ ∈ {0, 1} indicates the presence (1) or absence (0) of the invasive species from island *m*. For example, in a five island archipelago, a state where the first and fourth islands are invaded would be represented as *S_i_* = {1, 0, 0, 1, 0}. This definition of the archipelago’s invasion state follows standard metapopulation theory by focusing on the binary occupancy state of each island, but we admit that it does not include the size of the invasive population.

To define the probability of the archipelago transitioning between each system state, we need to construct a set of probabilistic transition matrices **T**(*k*). The elements of these matrices [**T**(*k*)]_*a,b*_ describe the probability that an archipelago in state *S_a_* this year, will transition to state *S_b_* next year. Management actions affect the metapopulation state, and the manager has *n* + 1 possible actions: attempting to eradicate invasive species from each of the *n* islands, and doing nothing (action *n* + 1).

The transition matrices can be constructed by analytically generating and then combining auxiliary extinction and colonisation matrices (Day and Possingham, 1995), but this method becomes complicated and computationally expensive for metapopulations as large as most invaded archipelagos (i.e., > 10 islands). We therefore approximate **T**(*k*) using Monte Carlo simulations of the SPOM (Bode and Possingham, 2007). After a large number of iterations, the mean of these matrices closely approximates the transition matrices.

### 2.2 Application of SDP

Armed with an archipelago SPOM, we can determine the best management strategy by applying SDP, a technique from operations research that has been applied to various problems involving decision-making under uncertainty (Dreyfus and Bellman, 1962). SDP creates a state-dependent optimal policy that identifies which island should be targeted for eradication, given that the archipelago is in a particular invasion state *S_i_*.

SDP first requires that a present conservation value *ψ*(*i*) be assigned to each of the archipelago states *S_i_*, indicating its value to managers (or other relevant stakeholders). We will assume that managers aim to eradicate invasives from as much of the archipelago area as possible. As a result, we let *ψ*(*i*) be the total area of the islands that are not occupied by the invasive species in state *S_i_*. The goal of the managers will be to maximise the expected net present conservation value of the archipelago, by choosing which island to target with an eradication project. Each action *k* carries a cost, *c*(*k*), and the system dynamics are determined by the transition matrix **T**(*k*).

Let *V* (*i, t*) denote the expected return on optimal actions after *t* years, when the problem starts in state *S_i_*. The optimal strategy is determined by applying the recursive, back-stepping dynamic programming equation:

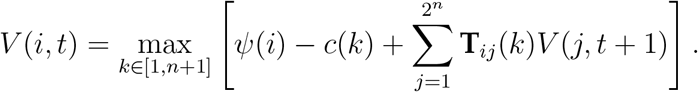

The recursion equation requires that a terminal value be defined, which we assume is the terminal state of the system *V* (*i, T*) = *ψ*(*i*). Starting at the terminal time, the managers identify the action *k* that maximises the expected value of the system in the penultimate step *T* − 1, in each system state *i*. This value is stored in the optimal policy matrix *A*(*i, T* − 1). The expected value of being in state *i* in any time step is the value of that state (*ψ*(*i*)), plus the expected value in the next time step (calculated using the appropriate transition matrix), minus the cost of taking the action *c*(*k*). This process is then repeated until the present time *t* = 1.

### 2.3 Case studies: the Seaforth and Dampier Archipelago

To illustrate the application of the SPOM/SDP framework, we consider two case studies: the eradication of rodents from the Seaforth archipelago in New Zealand, and the eradication of cane toads from the Dampier archipelago in Western Australia (Figure 1). The goal of this paper is not to offer prescriptive advice to ongoing eradication projects, and so our case studies are purposefully hypothetical: cane toads have not even reached the vicinity of the Dampier archipelago yet (although they will in the next few years; (Southwell et al., 2017)), and the Seaforth archipelago is located in Fiordland, where relatively few islands have been invaded by rats (Carter et al., 2021). These case studies illustrate the process of parameterising the decision-support tool, and are used to exemplify and analyse the results in a realistic setting.

**Figure 1:**
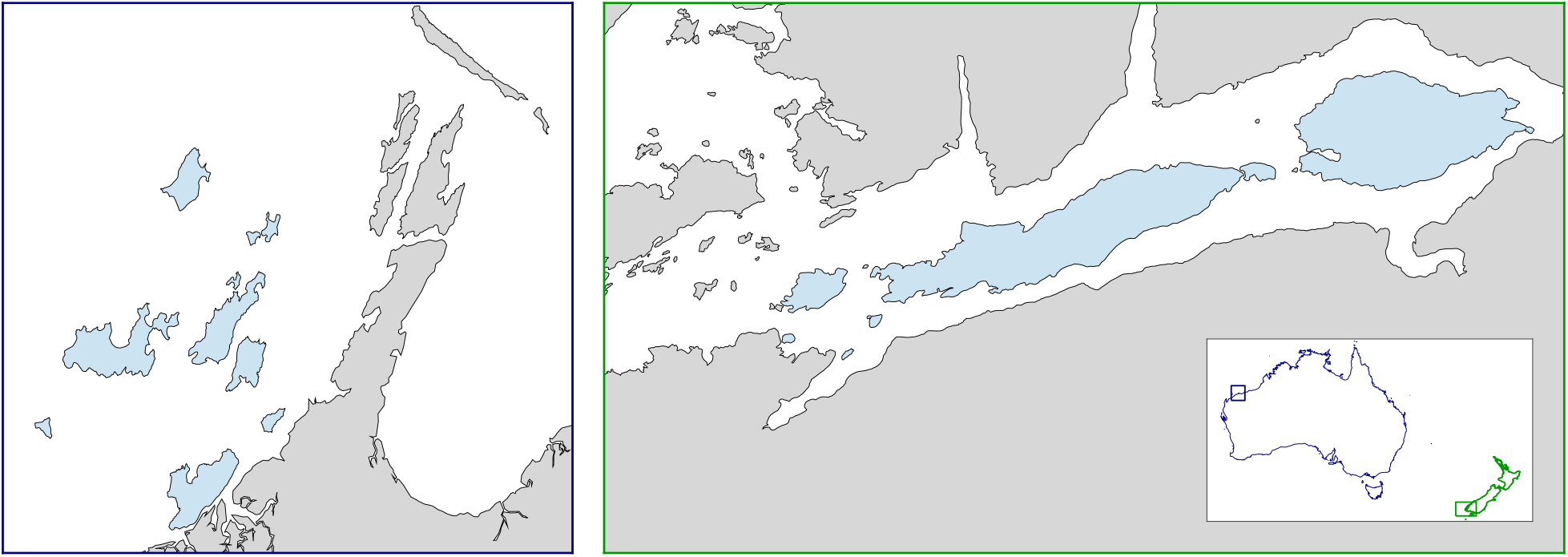
Maps of the locations of the (a) Dampier and (b) Seaforth case studies. Blue shaded islands indicate the archipelago being considered. Inset plot shows the location of the case studies in Australia (blue) and New Zealand (green).

For the Seaforth archipelago, we estimate the probability that rodents from island *a* will recolonise island *b* to be *p_ab_* = 0.2exp[−2.6*d*(*a, b*)], where *d*(*a, b*) is the shortest distance between the two islands in metres. We also calculate the probability that each island will be recolonised from the mainland based on the same equation, but using the minimum distance of each island to the mainland. Furthermore, we assume that recolonisation is impossible beyond a distance of 1 km. This distance, and the values of 2.6 and 0.2, are based on estimates of ship rat *Rattus rattus* colonisation abilities provided in (Lohr et al., 2017). The resulting recolonisation matrix is shown in Figure 2(a). We assume that a natural extinction is unlikely for this species, and place the probability at *e_a_* = 0.05. Using data on previous *Rattus* species eradications from the DIISE, we estimate that the probability of eradicating the species from an island with area *X* is given by the logistic function ln(*p*(1 − *p*)^−1^) = 1.7 − 0.027*X*.

**Figure 2:**
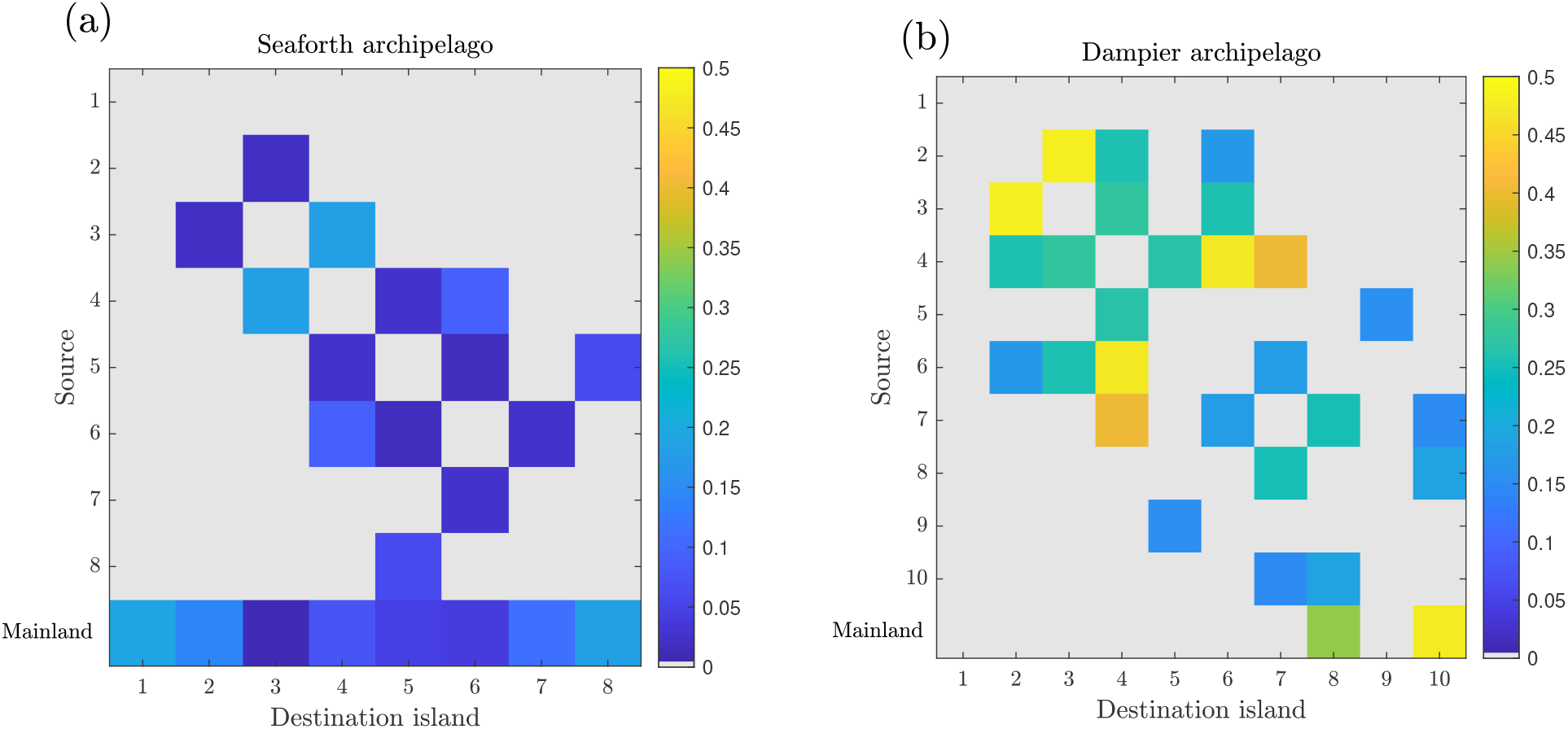
Heatmap visualisation of recolonisation matrices for (a) Seaforth and (b) Dampier case studies. Colours indicate the probability that an invaded row island (or mainland) would be able to reinvade a column islands. Seaforth islands are less likely to recolonise each other than Dampier islands. Seaforth islands are equally connected to the mainland, while Dampier islands have very different levels of mainland connection.

For the Dampier archipelago, we estimate the probability that cane toads from island *a*, or from the mainland, will colonise or recolonise island *b* as *p*(*a, b*) = 0.5exp[−0.00025*d*(*a, b*)], where *d*(*a, b*) is the shortest distance between two islands or the mainland. Recolonisation is assumed to be impossible beyond a distance of 5 km. The values are also based on estimates of colonisation abilities provided in (Lohr et al., 2017). The resulting recolonisation matrix is shown in Figure 2(b). We assume that a natural extinction is again unlikely for this species, and place the probability at *e_a_* = 0.05. We estimate the probability of successful eradication using estimates of cane toad eradication costs from (Smart et al., 2020). Assuming an annual eradication budget *B* of AUD $ 100, 000, and a fully-funded eradication cost per square kilometre of *D* = $96, 556, we set the probability of successful eradication of an island with area *α* to be *p*(*α*) = *B*(*αD*)^−1^.

Using the SPOM/SDP framework, we generate the optimal eradication policy for the Seaforth and Dampier archipelago. The initial state of the Seaforth archipelago case study consists of *n* = 8 islands, which are all initially occupied by rodents. There are *k* = 9 actions, representing a decision to target an island or to do nothing. Similarly, the initial state of the Dampier archipelago case study consists of 10 islands which are all occupied by cane toads, and there are 11 possible actions at each state. We allow the eradication projects to continue for twice the number of years required to eradicate throughout the archipelago, such that *T* = 16 years for the Seaforth archipelago, and *T* = 20 years for the Dampier archipelago. This ensures that there is enough time for a full eradication, given the possibility of setbacks from recolonisation and eradication failures.

### 2.4 The cost of ignoring recolonisation

We seek to measure the cost of ignoring reinvasion risk in island eradication prioritisations. We begin by calculating the optimal eradication strategy in the face of recolonisation risk. We then calculate the optimal eradication strategy under the assumption that recolonisation *cannot* occur (as assumed by current prioritisation tools). We then compare the performance of the “recolonisation” and the “no recolonisation” strategies, when they are applied to eradication projects where recolonisation *can* occur. We simulate both strategies on the Seaforth and the Dampier eradication problems, using 10,000 Monte Carlo simulations, beginning in the state where all islands are invaded, and running for *T* = 2*n* years. In this comparison, it is clear that the “recolonisation” strategy will outperform the “no recolonisation” strategy; our goal is to estimate the magnitude of the benefit.

## 3 Results

Applying SDP yields an optimal policy: a matrix state-dependent decision *A*(*i, t*) that maximises the expected conservation value of the archipelago. The optimal policy for a 10 island archipelago, over 20 years is contained in a 1024 × 20 matrix, a shape that makes visualisation challenging (see Supporting Figure 6). The optimal policy also changes through time, making it hard to intuit why particular islands were prioritised in particular states. Fortunately, the decisions recommended by the optimal policy can be readily mapped onto the archipelagos themselves.

Figure 3 shows the optimal sequence of eradications proposed for the Seaforth archipelago case study, when recolonisation can occur from either the mainland or other invaded islands. The green lines and dates in the figure represent the islands and dates that should be eradicated under the optimal policy, starting in 2026. The optimal eradication schedule tends to move from the east of the archipelago to the west, but not in a straightforward, linear manner.

**Figure 3:**
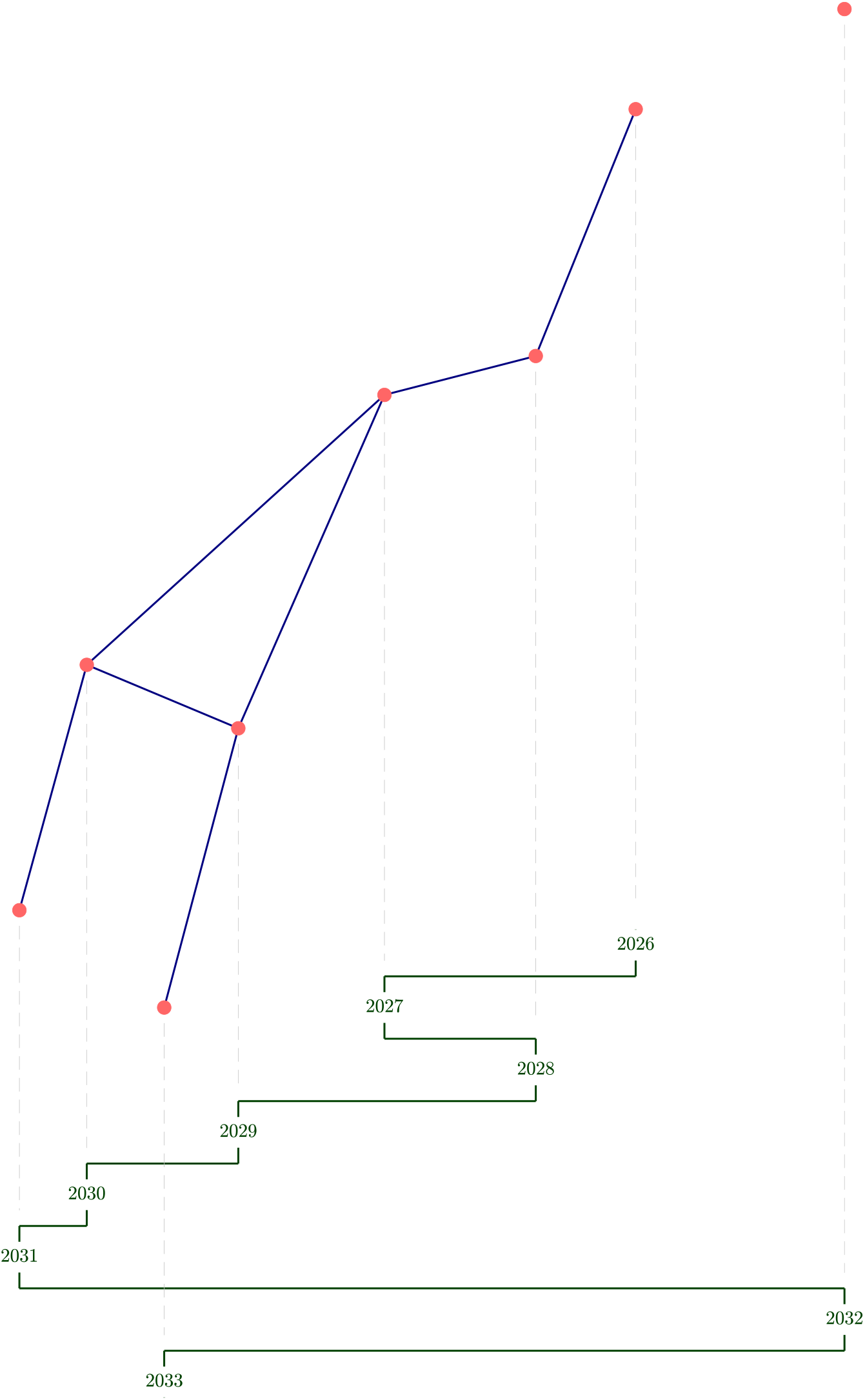
Results for the Seaforth archipelago example. Nodes show the islands in the archipelago in their approximate position, and edges show potential colonisation routes between the islands. All nodes are close enough to the south island of New Zealand to be recolonised by the mainland rat populations. Green lines and dates show the sequence of the optimal eradication, starting in 2026.

Figure 4 presents the optimal eradication sequence proposed for the Dampier archipelago case study with colonisation. In this case, the schedule of eradications begins with the most peripheral islands in the archipelago, with a preference for islands that can be recolonised by the fewest other islands. The last islands in the sequence are those closest to mainland populations of the invasive species.

**Figure 4:**
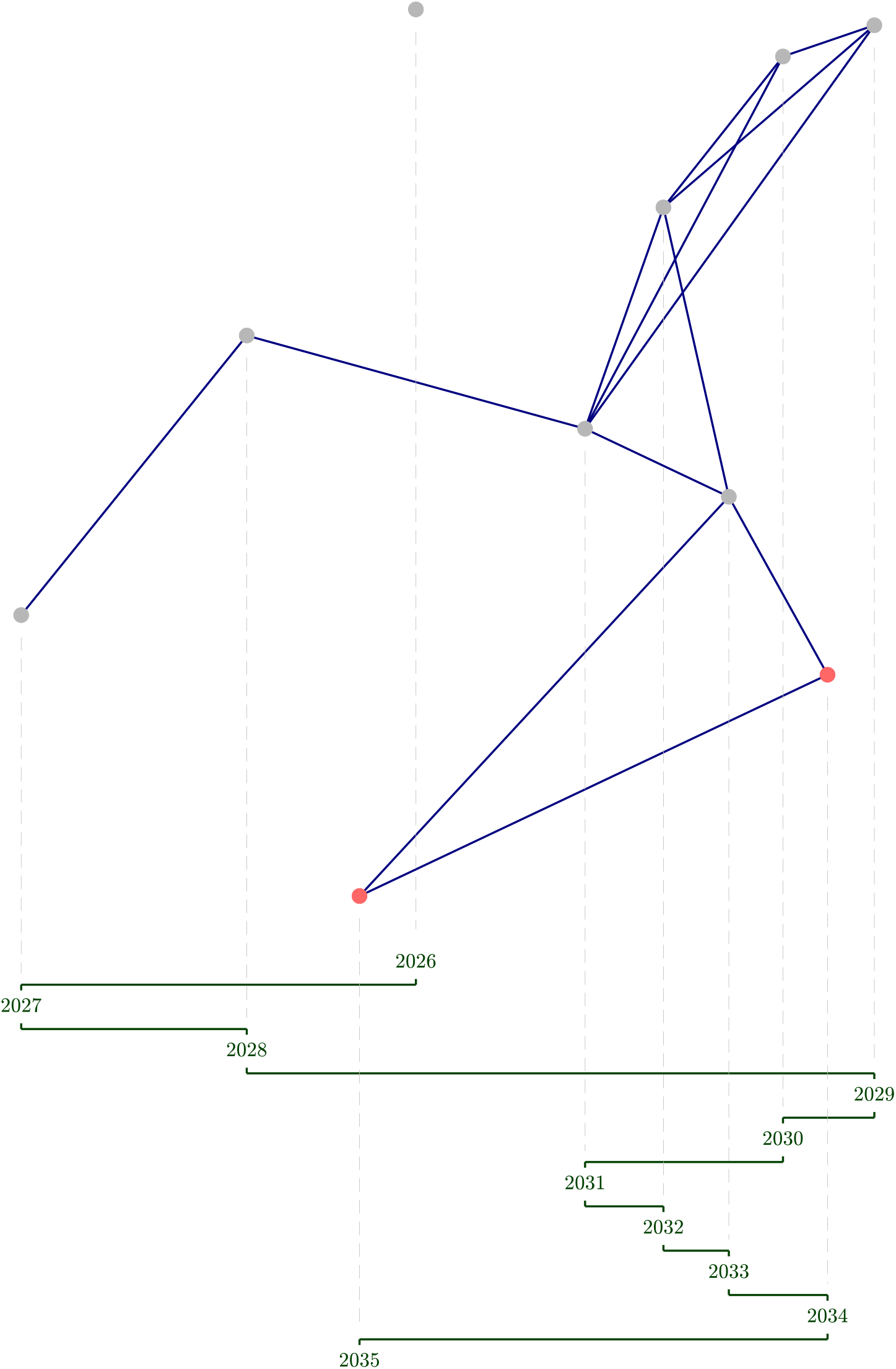
Results for the Dampier archipelago example. Nodes show the islands in the archipelago in their approximate position, and edges show potential colonisation routes between the islands. Nodes with red shading are close enough to the Australian mainland that they can be recolonised by mainland cane toad populations. Nodes with grey shading cannot be directly reached by cane toads from the mainland. Green lines and dates show the sequence of the optimal eradication, starting in 2026.

For both archipelagos, the inclusion of reinvasion risk completely altered the optimal eradication sequence (compare Figures 3 and 4 with Supporting Figures 9 and 10). In general, the optimal solution responded to the possibility of reinvasion by taking a more spatially autocorrelated approach to eradication, targeting islands one-by-one along the backbone of an archipelago.

The benefits of considering recolonisations varied between the two example archipelagos. Figure 5 shows the distribution of the expected conservation benefits for both the “recolonisation” and “no recolonisation” optimal policies. Including recolonisation in the analyses improves outcomes for both the Seaforth and Dampier archipelago, but by notably different amounts. In the Dampier archipelago, including recolonisation delivers outcomes that are 42% better (on average) than omitting recolonisation. For the Seaforth archipelago by contrast, the improvement is only 6% on average.When recolonisation dynamics are included in the prioritisation process, the optimal state-dependent policy changes for 33% of decisions in the Seaforth archipelago (Supporting Figure 8), and for 72% of decisions in the Dampier archipelago (Supporting Figure 7).

**Figure 5:**
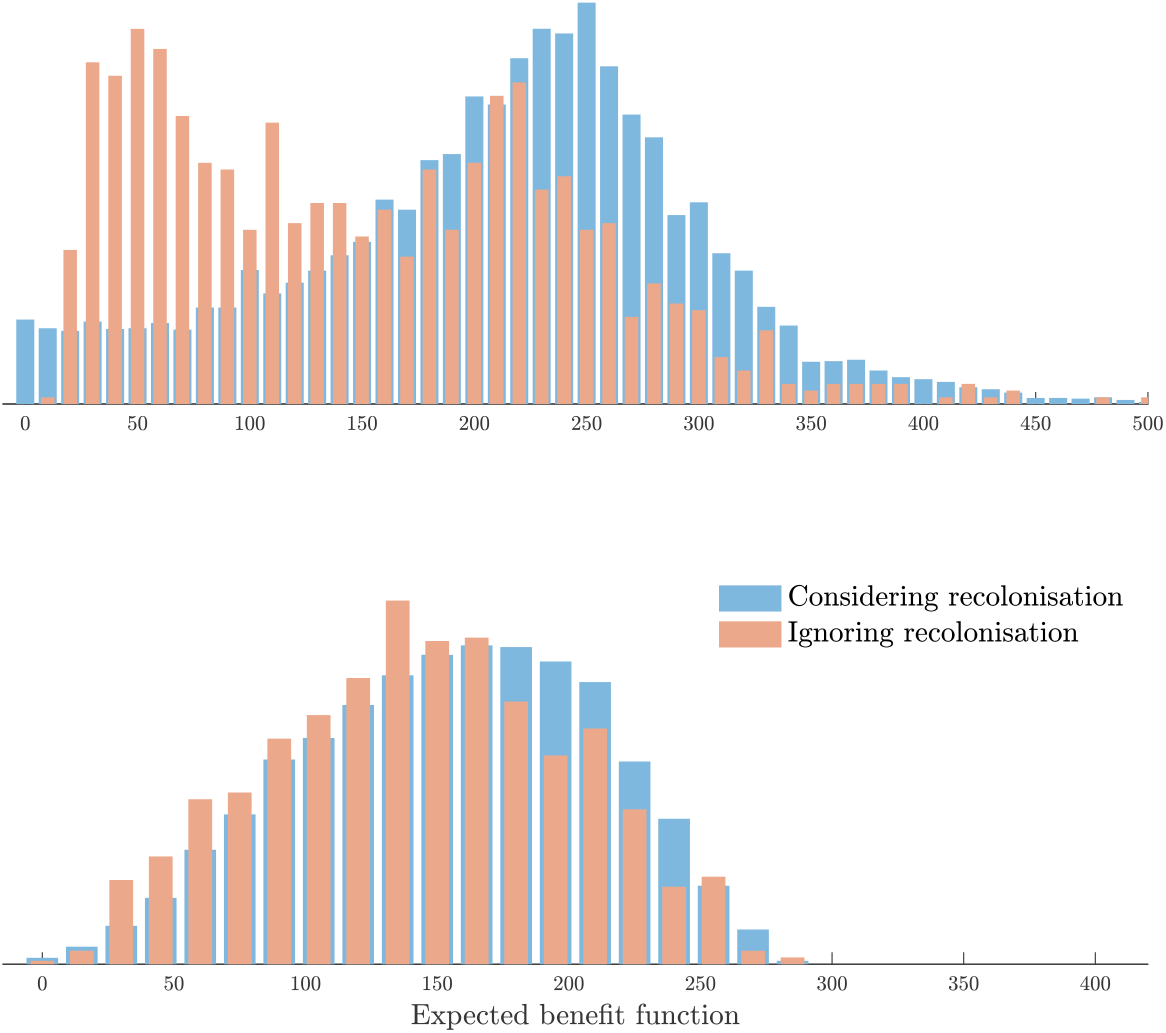
Performance of the optimal solution where recolonisation is considered (shaded blue), and where recolonisation is not considered (shaded brown). The upper panel portrays the results for the Dampier archipelago. Including recolonisation improves the expected return by 42 %. The lower panel shows the results for the Seaforth archipelago. Including recolonisation improves the expected return by 6 %.

## 4 Discussion

By combining ideas from standard island eradication prioritisation methods and metapopulation theory, we have created a decision-support tool that considers both the intrinsic qualities of each island in an archipelago, and the system-wide and state-dependent dynamics of recolonisation. Our analyses show how prioritisations can be calculated for two hypothetical multi-island eradication projects (Figures 3 & 4), and reveal the limitations of a “list-based” approach to eradication planning. Static priority lists assume that rankings do not change as action is taken. Given the frequency with which invasive species can colonise unoccupied islands, only a state-based approach to prioritisation can claim to offer efficient outcomes.

Reinvasion risk impacts both the choice of which island to eradicate invasive species from, and also the outcomes of those eradication choices. Of course, the importance of considering reinvasion risk varies with the likelihood that reinvasion will occur. In our examples, the inclusion of reinvasion risk was more impactful in the Dampier archipelago, although its effects on decisions and outcomes in the Seaforth archipelago were not negligible. The disparity between the two examples can be attributed to differences in the invasive species and the archipelagos being invaded. The islands of the Seaforth archipelago are less connected to each other than those in the Dampier archipelago, and more uniformly connected to the mainland (Figure 2). Both factors make recolonisation a less influential factor in the Seaforth archipelago. Weaker connections between the islands makes recolonisation a less important process, and make a “no recolonisation” assumption closer to the truth. By contrast, when connections to the mainland are more heterogeneous in strength, as in the Dampier archipelago, it becomes more important to understand which islands are acting as a conduit for recolonisation.

Our approach has three primary limitations. First, the population dynamics modelled by SPOMs are relatively simple: islands can only be eradicated or invaded. Although this is a common assumption in between-island prioritisation models (Baker and Bode, 2021), it neglects the effects of abundance on the challenges of eradication, or on the probability that a population will act as a source of recolonisation. Similarly, the actions available at each time step in the model are limited, only allowing a single island to be targeted for eradication each year (or none), and only considering a single eradication method (Baker et al., 2017).

Second, our use of SDP places computational limits on the archipelagos that can be analysed. Here, we consider an 8 and 10 patch model; however, for a larger archipelago the number of system states would soon become unmanageable. We estimate that an archipelago consisting of 15 or more islands would be impossible without approximation methods (Nicol et al., 2010).

Finally, we have not considered observation error. Our analysis is underpinned by the assumption that at all times, managers are aware of the invasion state of each island in the archipelago, even when they have only recently been (re)invaded. While this is a common assumption in island eradication prioritisation, it is quite unrealistic. Managers must often wait long periods prior to determining the success of an eradication (Ramsey et al., 2009), and reinvasion populations are initially small and cryptic. Island prioritisation theory is in dire need of decision-support tools to address the issue of invasion state uncertainty, particularly when reinvasion processes cause the system state to change continually.

Island eradications have delivered enormous and lasting benefits for threatened species and ecosystem conservation throughout the world (Jones et al., 2016). However, their efficacy is jeopardised by the risk of reinvasion, which not only undermines the benefits accruing to each island, but also the rationale for island eradications as a strategy. Our results emphasise both the dangers posed by reinvasion, and the benefits of incorporating reinvasion risk in eradication prioritisation methods.

## Author contributions

All authors conceived the research idea. All authors contributed to the methods, writing and editing. VK, AN, and MB contributed additional editing and writing.

## Acknowledgements

The authors would like to acknowledge valuable discussions with Geoffrey Wong, Louis Dweck, Jaymie Tilbury and Sharafat Hassani on the methods used in this paper.

## Appendix

MATLAB code used to generate the figures presented in this paper is available at https://github.com/vineyk24/Island-Eradication.

## Supporting Information

**Figure 6:**
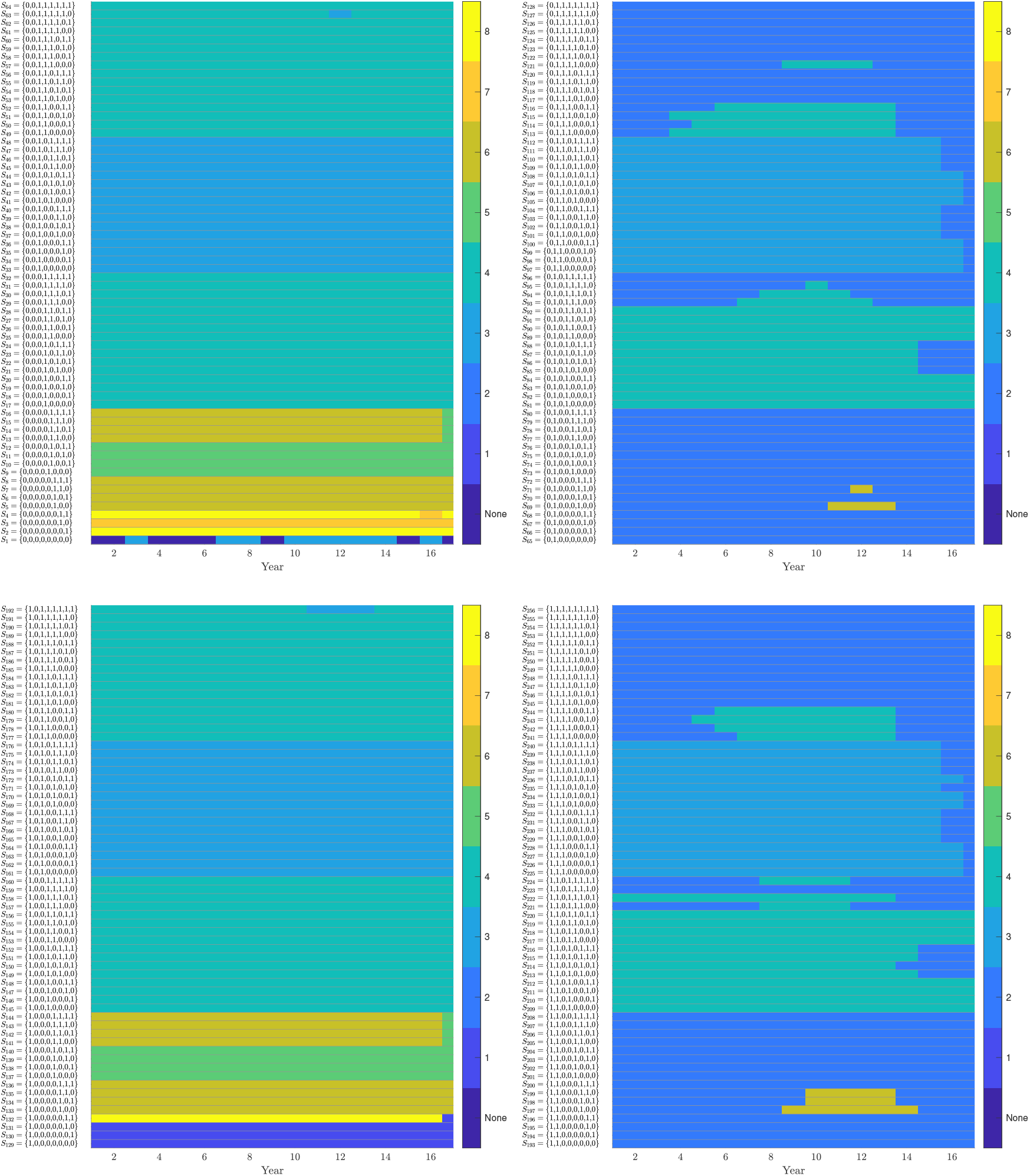
Optimal eradication policy for the Seaforth archipelago example. Colours show the optimal island to target for eradication in a given year (columns), in a given state (rows). Policy has been broken into four sections for labelling and readability.

**Figure 7:**
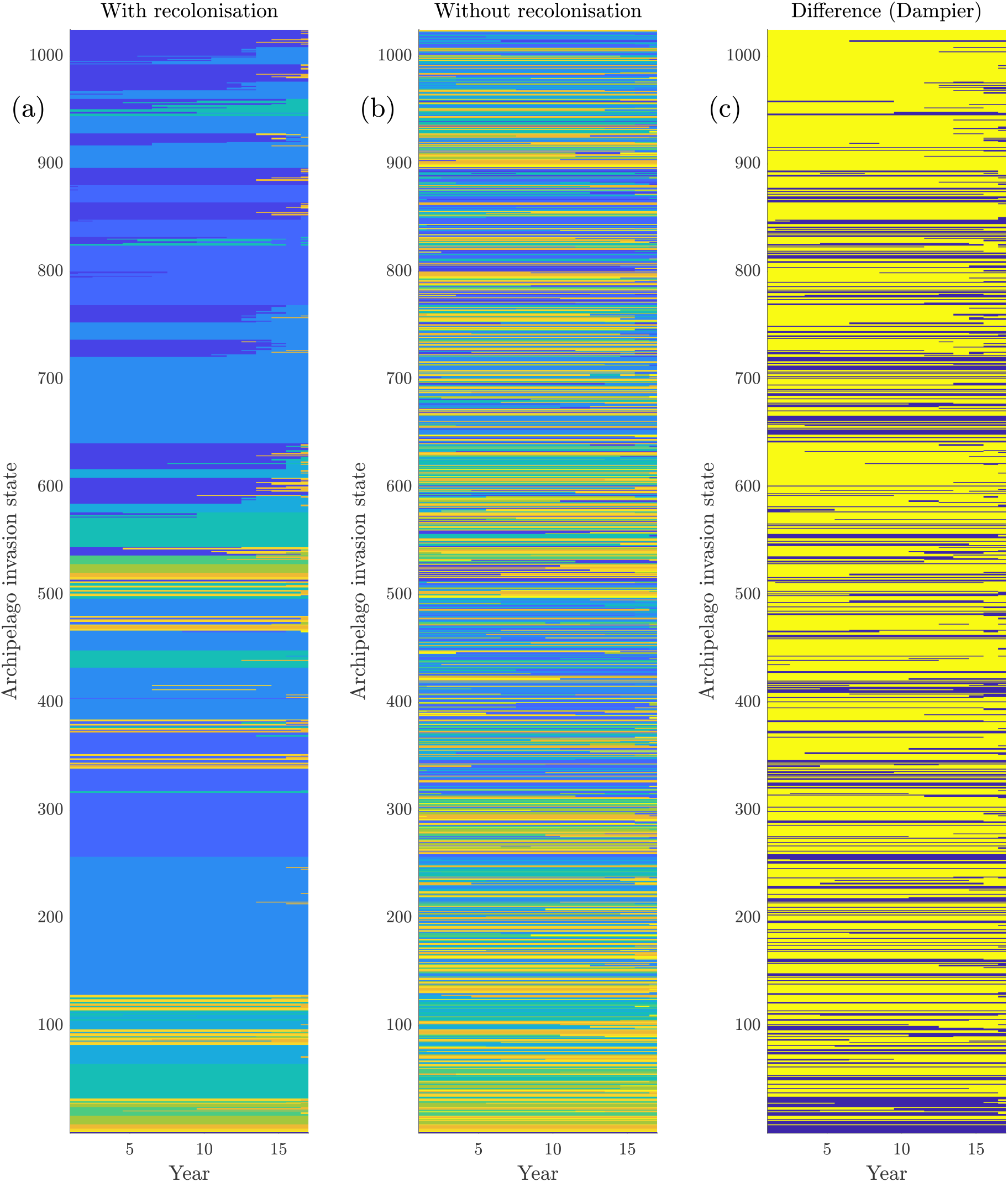
Optimal eradication policy for the Dampier archipelago case-study, with and without recolonisation being considered. The purpose of this figure is not to communicate the optimal strategy under the two assumptions, but just to communicate the degree of agreement between their optimal solutions. In panels (a) and (b), colours indicate the optimal island to target for eradication in a given year (columns), in a given state (rows). Colours carry the same meaning as in Figure 6. Panel (c) indicates the differences between panels (a) and (b), with yellow indicating disagreement, and blue indicating agreement. In the Dampier case-study, it’s clear that the two solutions are very different.

**Figure 8:**
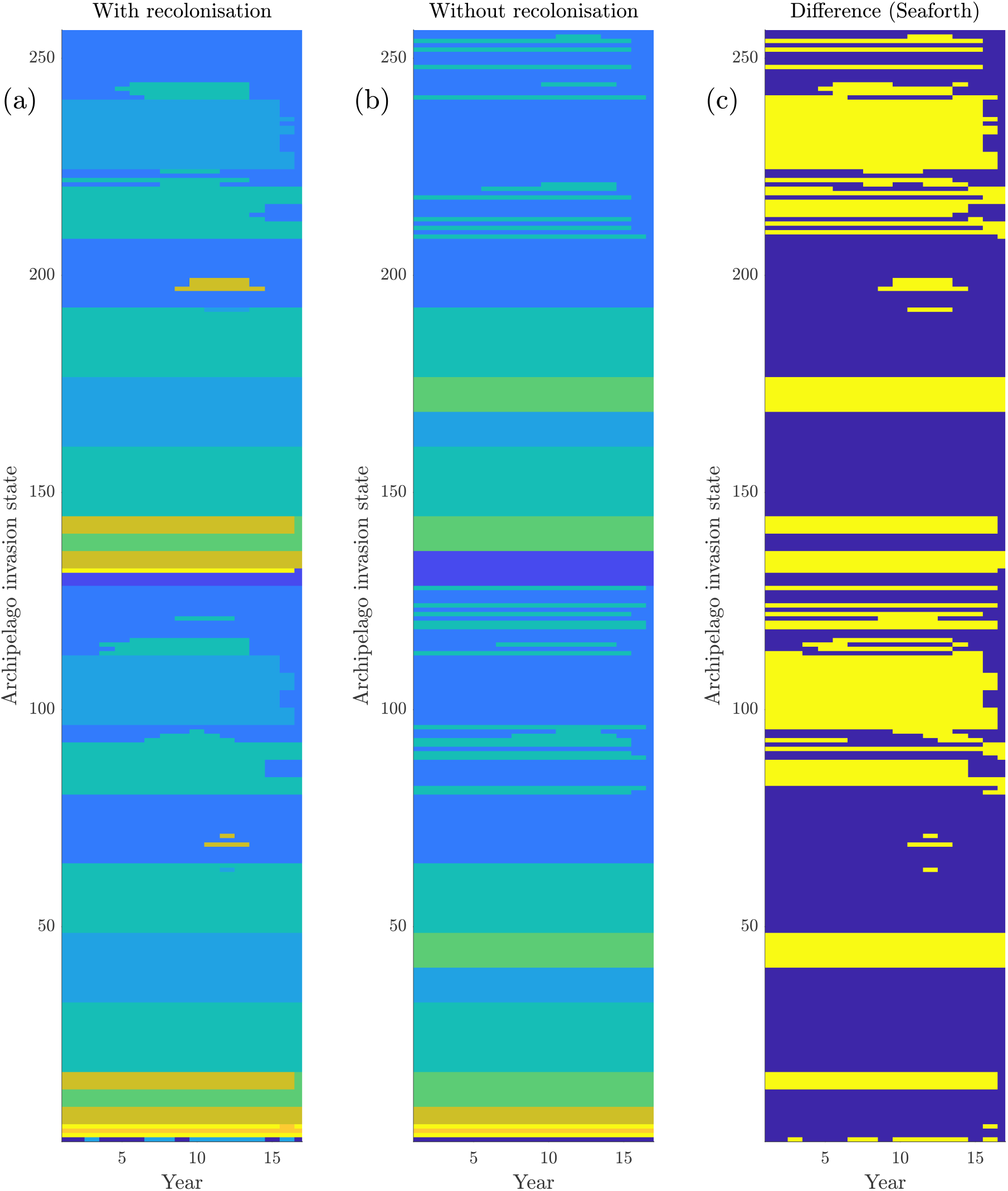
Optimal eradication policy for the Seaforth archipelago case-study, with and without recolonisation being considered. The purpose of this figure is not to communicate the optimal strategy under the two assumptions, but just to communicate the degree of agreement between their optimal solutions. In panels (a) and (b), colours indicate the optimal island to target for eradication in a given year (columns), in a given state (rows). Colours carry the same meaning as in Figure 6. Panel (c) indicates the differences between panels (a) and (b), with yellow indicating disagreement, and blue indicating agreement. There is more agreement between the optimal solutions in the Seaforth case-study, than in the Dampier case-study.

**Figure 9:**
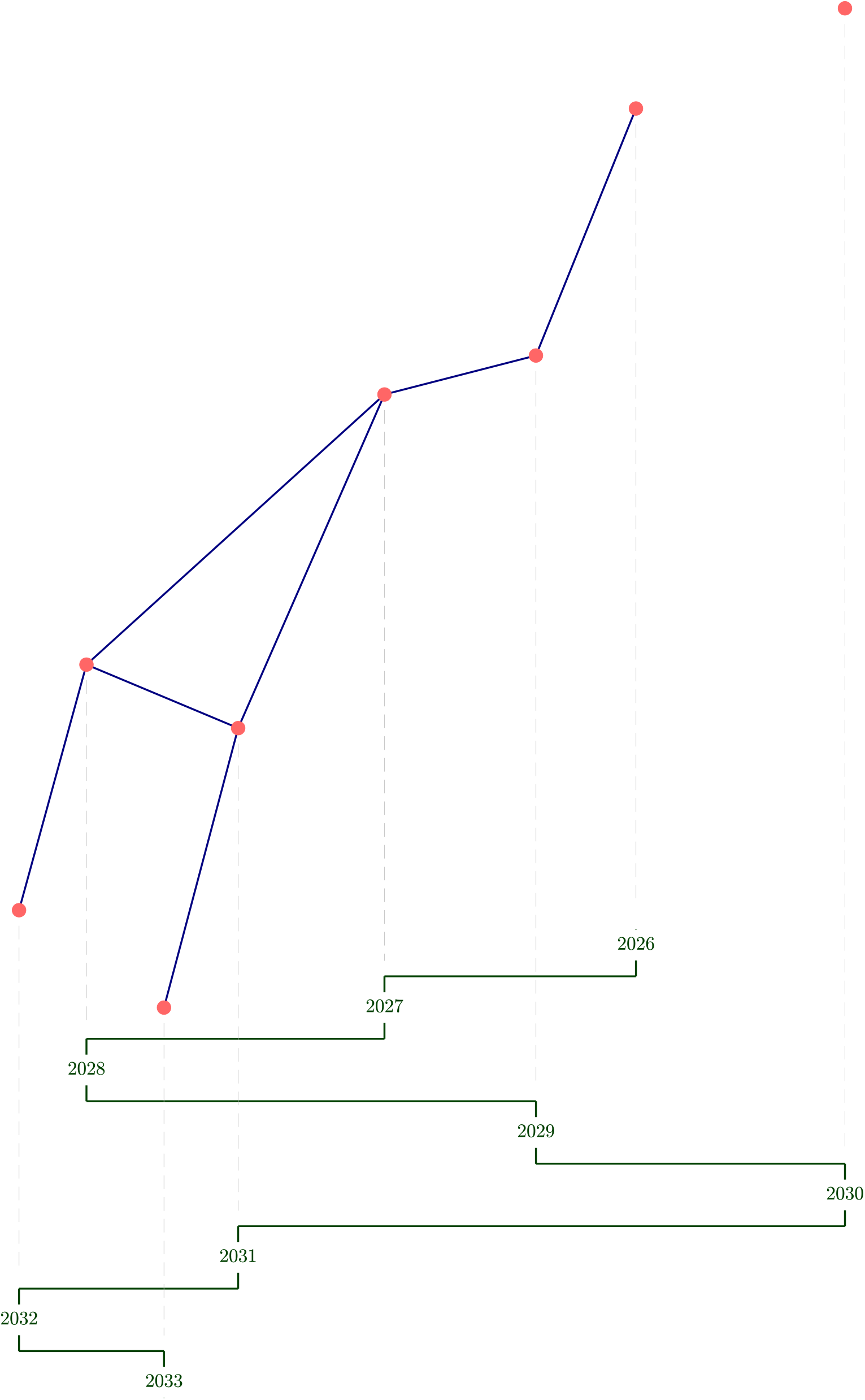
Results for the Seaforth archipelago example, when recolonisation is not included. The purpose of this figure is to contrast with Fig 3, where recolonisation is included. Nodes show the islands in the archipelago in their approximate position, and edges show potential colonisation routes between the islands (although these are not operating in this version of the model). All nodes are close enough to the south island of New Zealand to be recolonised by the mainland rat populations, although again, this process is not operating here. Green lines and dates show the sequence of the optimal eradication, starting in 2026.

**Figure 10:**
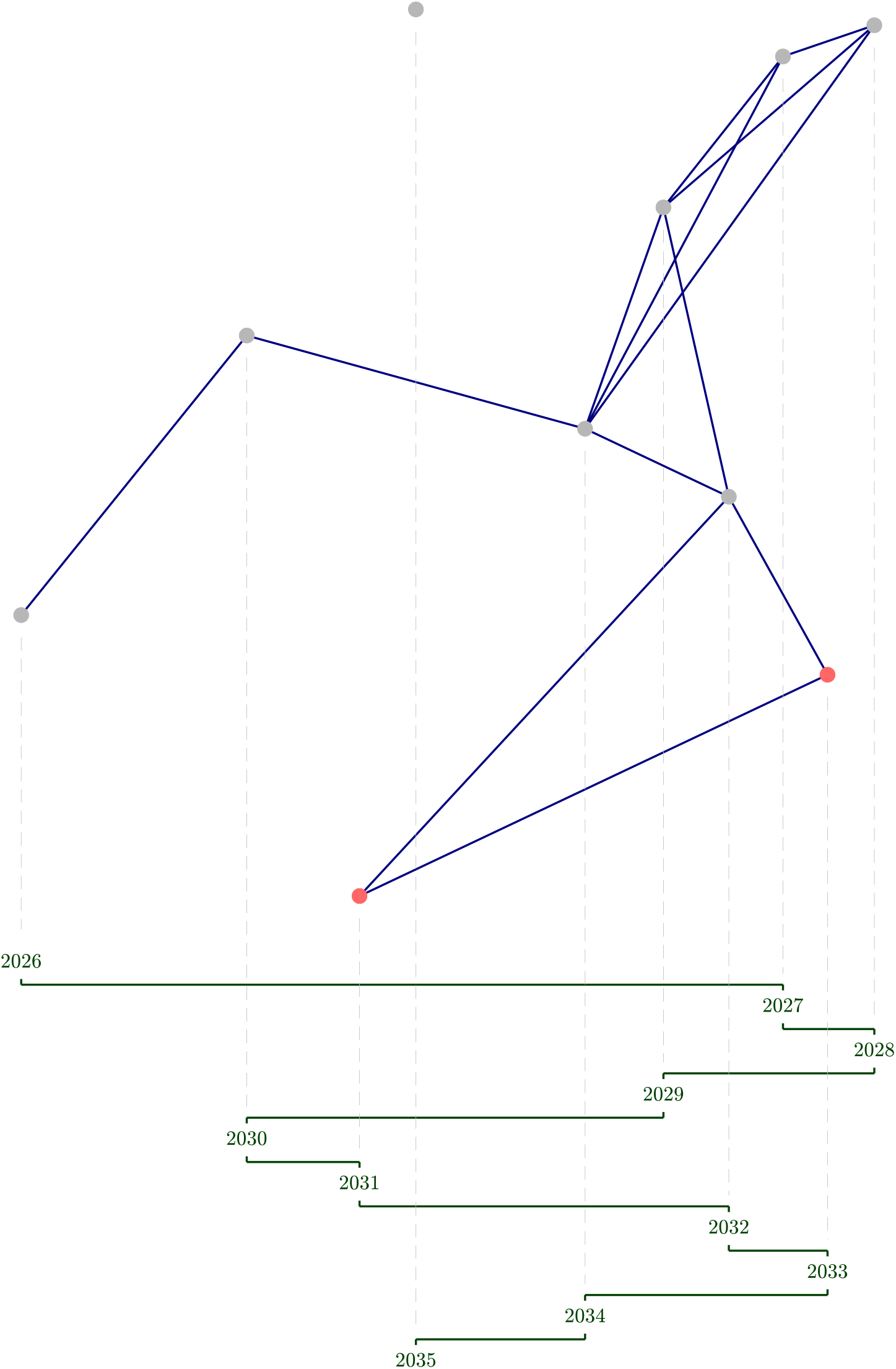
Results for the Dampier archipelago example, when recolonisation is not included. The purpose of this figure is to contrast with Fig 4, where recolonisation is included. Nodes show the islands in the archipelago in their approximate position, and edges show potential colonisation routes between the islands (although these are not operating in this version of the model). Green lines and dates show the sequence of the optimal eradication, starting in 2026.

